# pylifemap: Mapping Data onto the Tree of Life

**DOI:** 10.64898/2025.12.16.694609

**Authors:** Julien Barnier, Cassandra Bompard, Aurélie Siberchicot, Vincent Navratil, Joaquim Martin, Damien M. de Vienne

**Affiliations:** Université de Lyon, Université Lyon 1, UMR CNRS 5558 Laboratoire de Biométrie et Biologie Évolutive, 69622 Villeurbanne, France; PRABI, Pôle Rhône-Alpes Bioinformatics Center, Université Claude Bernard Lyon 1, 69622, Villeurbanne, France; Institut Français de Bioinformatique, IFB-Core, UMS 3601, 2 rue Gaston Crémieu, Évry Cedex 91057, France; European Virus Bioinformatics Center, Leutragraben 1, Jena 07743, Germany; Independent Software Engineer, 38200 Saint-Sorlin-de-Vienne, France; Biodiversity Computing Group, Institute of Computer Science, Foundation for Research and Technology Hellas, 100 Nikolaou Plastira, 70013 Heraklion, Crete, Greece

**Author notes:** Corresponding authors: DAMIEN M. DE VIENNE, JULIEN BARNIER, Université Lyon 1, CNRS, Laboratoire de Biométrie et Biologie Évolutive, Bâtiment Mendel 43 boulevard du 11 Novembre 1918, 69622 VILLEURBANNE CEDEX, **E-mails:**.

**Keywords:** Lifemap, metagenomics, taxonomic classification, visualization, Tree of Life

## Abstract

The need to visualize data associated with NCBI Taxonomy Identifiers is growing in various biological fields ranging from comparative genomics to metagenomics and metabarcoding, and even for outreach.
No tool today allows visualization of such data while still keeping the full vision of the whole taxonomy, possibly causing a biased view of the data at hand.
Here we introduce *pylifemap*, a Python package that allows users to map their own data directly on the interactive taxonomic tree proposed by Lifemap. Through a simple syntax, layers (points, lines, icons, heatmaps, etc.) depicting any type of data are superimposed on the Lifemap basemap, producing an interactive map to inspect biological datasets. The produced visualizations can easily be shared with others through notebooks or standalone HTML files, or exported as static images.
We illustrate the utility of *pylifemap* in the exploration of two contrasting datasets: the IUCN Red List of Threatened Species and the output of a large-scale environmental metagenomics experiment.

**Data/Code for peer review:** *pylifemap* is available at PyPI and easily installable with pip or uv. The development version can be found at https://lifemap-tol.github.io/pylifemap/ along with extensive documentation and examples.

## 1. Introduction

Visualization of large datasets associated with taxonomic identifiers is increasingly essential across various fields. In metagenomics, for instance, sequenced biological samples and their subsequent analysis lead to lists of taxa, identified by their taxonomic identifiers, associated with estimated abundance information. In comparative genomics, genomic data relevant to groups of species can be computed and analyzed. In ecology, diverse ecological data are routinely obtained for species, with their NCBI taxonomy identifiers (taxids) being extractable. In all these cases, as well as for outreach activities related to taxonomy, systematics, and the Tree of Life, visualizing such data is crucial because it facilitates accurate interpretation, helps reduce possible biases and supports the dissemination of knowledge.

In 2016, we released Lifemap (https://lifemap.cnrs.fr), a web application for the interactive exploration of the entire NCBI taxonomy (de Vienne 2016). The Lifemap explorer resembles modern interactive geographical maps, except that the map is taxonomical instead of geographical. Exploration of the full taxonomy (as provided by the NCBI, Federhen 2012; Schoch et al. 2020) is done by zooming and panning, and only the relevant information is given at each zoom level. This way, the entire taxonomy can be explored intuitively and interactively. Lifemap has become over the years a reference for exploring and presenting the Tree of Life with tens of thousands of visitors every month. In addition to being a useful resource to explore taxonomy and get easy access to related databases (e.g. NCBI, IUCN, GBIF 2025; iNaturalist 2025, etc.), Lifemap also provides a unique interactive map of the whole taxonomy that can be reused for other purposes.

By employing Lifemap as a basemap, we propose here a tool, *pylifemap*, for easily displaying information related to various taxa on the map directly. With this Python package, users can input and visualize data of any type associated with taxids. These identifiers are used to locate the position of each taxon on the Lifemap basemap, and user-defined layers (points, lines, heatmaps, etc.) are overlaid on the map to display the desired information. These layers can represent different aspects of the data, and users can define as many layers as necessary to suit their specific needs.

In order to illustrate the use of *pylifemap* and briefly present its underlying syntax, we used two datasets. The first one is the IUCN Red List of Threatened Species, a list of 172,620 animals, fungi and plants categorized based on their risk of extinction (IUCN 2025). The second is the output of the taxonomic classification tool Kraken2 (Wood et al. 2019) applied to environmental samples collected in 2020 in the Huanan seafood market in Wuhan, China (Liu et al. 2024). To facilitate comparison and visualization, this dataset was combined with another one listing the wild animal species sold in the same market just prior to the pandemic (Xiao et al. 2021). This comparison is important for understanding the potential origins of the SARS-CoV-2 pandemic (Crits-Christoph et al. 2024).

As the volume of available sequence data continues to grow (e.g. Katz et al. 2022), downstream analyses will inevitably generate an increasing number of datasets as well. Approaches such as the one proposed here for visualizing extensive data on an interactive and exhaustive taxonomic map may soon become a standard practice.

## 2. *pylifemap* overview

*pylifemap* allows mapping data associated with NCBI taxids on the interactive taxonomy explorer Lifemap (de Vienne 2016). To do so, *pylifemap* takes as input a pandas or Polars DataFrame with one column listing the taxids of interest and other columns containing associated data of any kind (numerical, categorical, textual, etc.). From this, *pylifemap* proceeds through a two-step process: first, it calls the Lifemap database to retrieve the coordinates of the different taxids and of their ancestors on the map. Second, it maps the data provided by the user at the correct location on the map, following the graphical and mapping instructions given by the user.

The basemap (or background map) on which the data are plotted is the same as the one on which the public Lifemap website relies (https://lifemap.cnrs.fr), although it can be styled differently for cleaner visualization and easier integration of the produced visualizations in publications. The taxonomy displayed in the basemap is automatically updated weekly, in order to always be up to date with the latest version of the NCBI taxonomy (Federhen 2012; Schoch et al. 2020).

Under the hood, the visualizations produced by *pylifemap* are Jupyter widgets powered by the *anywidget* framework (Manz et al. 2024), which enables integration of Python code with interactive JavaScript visualizations. These widgets can be produced and used interactively in notebook-like environments such as Jupyter, Marimo or Quarto, or generated from a Python script to be viewed in a web browser or saved as an HTML file.

### Visualization options and basics syntax

*pylifemap* proceeds by the superposition of layers on the basemap. Each layer can display either points, lines, text, icons, heatmap, donut charts or screengrid, and graphical options allow these layers to be precisely parametrized (e.g. colors, radius size, line width, etc., see Table 1A) according to the data. Other graphical elements can also be defined (Table 1B), including popups (opening on click) that can be parametrized by the user and associated with each element in each layer.

**Table 1.**
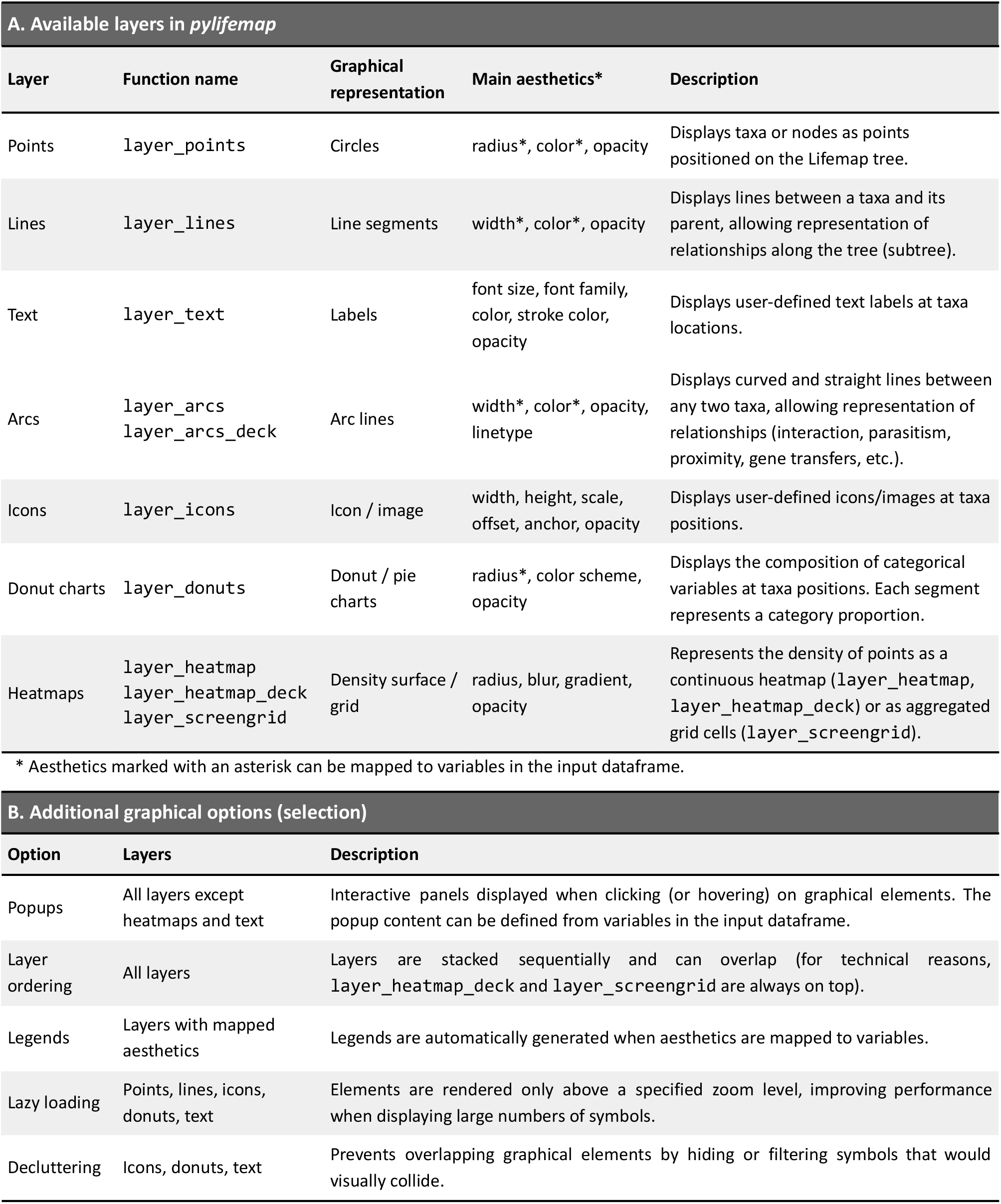
Visualization layers (A) and main additional graphical options (B) available in *pylifemap* (v 0.2.0).

| A. Available layers in *pylifemap* |  |  |  |  |
| --- | --- | --- | --- | --- |
| Layer | Function name | Graphical representation | Main aesthetics* | Description |
| Points | `layer_points` | Circles | radius*, color*, opacity | Displays taxa or nodes as points positioned on the Lifemap tree. |
| Lines | `layer_lines` | Line segments | width*, color*, opacity | Displays lines between a taxa and its parent, allowing representation of relationships along the tree (subtree). |
| Text | `layer_text` | Labels | font size, font family, color, stroke color, opacity | Displays user-defined text labels at taxa locations. |
| Arcs | `layer_arcs`<br>`layer_arcs_deck` | Arc lines | width*, color*, opacity, linetype | Displays curved and straight lines between any two taxa, allowing representation of relationships (interaction, parasitism, proximity, gene transfers, etc.). |
| Icons | `layer_icons` | Icon / image | width, height, scale, offset, anchor, opacity | Displays user-defined icons/images at taxa positions. |
| Donut charts | `layer_donuts` | Donut / pie charts | radius*, color scheme, opacity | Displays the composition of categorical variables at taxa positions. Each segment represents a category proportion. |
| Heatmaps | `layer_heatmap`<br>`layer_heatmap_deck`<br>`layer_screengrid` | Density surface / grid | radius, blur, gradient, opacity | Represents the density of points as a continuous heatmap (`layer_heatmap`, `layer_heatmap_deck`) or as aggregated grid cells (`layer_screengrid`). |
| * Aesthetics marked with an asterisk can be mapped to variables in the input dataframe. |  |  |  |  |
| B. Additional graphical options (selection) |  |  |  |  |
| Option | Layers | Description |  |  |
| Popups | All layers except heatmaps and text | Interactive panels displayed when clicking (or hovering) on graphical elements. The popup content can be defined from variables in the input dataframe. |  |  |
| Layer ordering | All layers | Layers are stacked sequentially and can overlap (for technical reasons, `layer_heatmap_deck` and `layer_screengrid` are always on top). |  |  |
| Legends | Layers with mapped aesthetics | Legends are automatically generated when aesthetics are mapped to variables. |  |  |
| Lazy loading | Points, lines, icons, donuts, text | Elements are rendered only above a specified zoom level, improving performance when displaying large numbers of symbols. |  |  |
| Decluttering | Icons, donuts, text | Prevents overlapping graphical elements by hiding or filtering symbols that would visually collide. |  |  |

The syntax used by *pylifemap* for visualization is straightforward. A general function `Lifemap()` is used to load the dataset, and to define general parameters for the map such as the theme, the width and height, the initial center and zoom level, etc. Then a list of layer methods (Table 1A) are used to define the elements present on the map, their appearance and their dependence on values of the input dataframe (inherited from the `Lifemap()` function or defined for each layer). Finally, the `Lifemap.show()` and the `Lifemap.save()` functions are used to respectively open the map for interactive visualization (in the browser or as a widget) or save the map as an HTML file for easy sharing and integration in websites. As an example, for a dataframe `df` the basic syntax for plotting points for taxa in column `“taxid”` with radius proportional to column `“value”` is:

~~~
Lifemap(df).layer_points(radius=“value”).show()
~~~

More syntax examples are provided below along with the biological illustrations.

### Inferring values at parent nodes

*pylifemap* also provides functions to compute values associated with parent nodes in the taxonomy for each variable of the initial dataset. Three such “aggregation” functions are provided: `aggregate_count()` to count at each node the number of descendant taxids present in the initial dataset, `aggregate_num()` to apply a function (min, max, mean, median and sum) to numerical variables associated with taxids present in the dataset, and `aggregate_freq()` to compute frequencies of categorical variables associated with the taxids in the initial dataset.

These functions (and other ones, more technical and not detailed here) help visualizing very large datasets, because the information present at deep zoom levels in the taxonomy can be summarized at lower zoom levels (higher taxonomic ranks), leveraging the amount of information to be plotted simultaneously on the map.

### Utility functions

Utility functions are provided along *pylifemap* to better characterize the dataset at hand: the function `get_unknown_taxids()` lists the taxids in the dataset that are unknown in Lifemap (e.g. in the latest version of the NCBI taxonomy presented in Lifemap, i.e. one week old at most); the function `get_duplicated_taxids()` retrieves the list of taxids in the dataset that are duplicated. Note that only the first occurrence of the taxid in the dataset (first row where it appears) is used for plotting in *pylifemap*. In case of unknown or duplicated taxids in the dataset, visualization functions still work but a warning is issued to the user.

## 3. Illustrative example of the use of *pylifemap* on biological datasets

In order to demonstrate how *pylifemap* can be utilized, we present here two different examples of its usage in the visualization of datasets in distinct biological contexts. Note that because the Lifemap explorer is interactive by nature, rendering the visual experience on static images is not ideal. The examples presented below with a single image export (Figure 1) are also available online for interactive exploration in a gallery (along with other examples) accessible at https://lifemap-tol.github.io/pylifemap/gallery/gallery.html.

**Figure 1.**
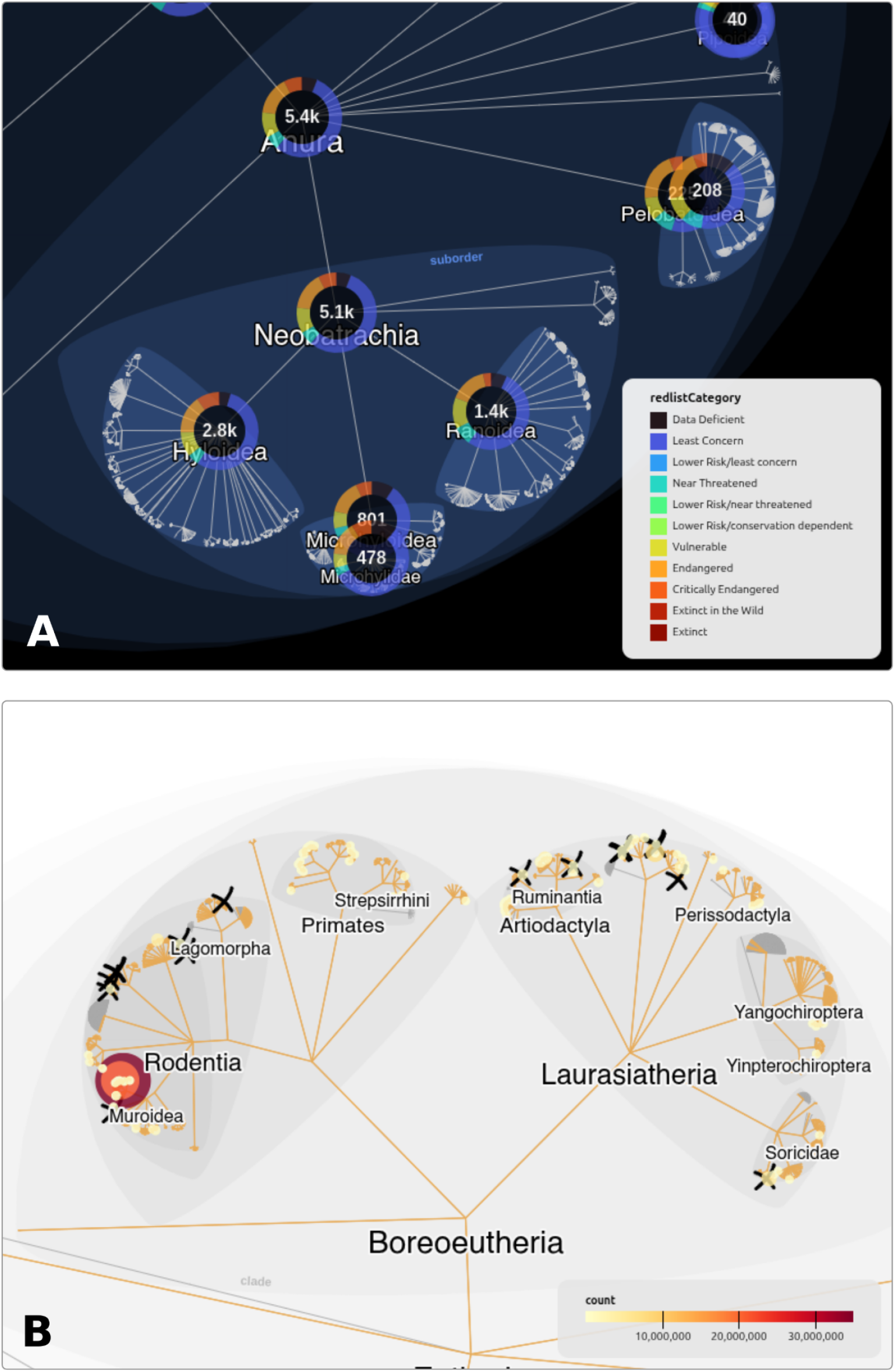
Two exports of interactive visualizations obtained with *pylifemap*. **A**.Visualization of the proportion of species at each taxonomic rank that belong to each one of the 11 IUCN Red List categories. Numbers inside donuts represent the total number of species belonging to each clade catalogued by IUCN (data from IUCN 2025). **B**. Complex visualization of the Wuhan market metagenomics datasets with three layers: (i) the number of reads mapping to taxa – as represented by the color and size of the points – following a taxonomic classification with Kraken2 (data from Liu et al. 2024, see text); (ii) the paths to the taxa available in the reference GenBank nt database used as a reference for Kraken2, represented by orange lines along the tree branches; (iii) the wild animals sold on the Wuhan market just before the pandemics, highlighted by black crosses (data from Xiao et al. 2021). The interactive visualizations associated with these snapshots are available in the *pylifemap* gallery: https://lifemap-tol.github.io/pylifemap/gallery/gallery.html.

### IUCN Red List

The International Union for Conservation of Nature’s Red List of Threatened Species is an indicator of the health of the world’s biodiversity. Focusing on animals, plants and fungi, it provides an up-to-date extinction risk status for 172,620 species (IUCN 2025).

For 103,116 of these species, we could recover their corresponding NCBI taxid by mapping their Latin name to the latest taxonomy using the ete3 Python package (Huerta-Cepas et al. 2016). We then used the function `aggregate_freq()` of *pylifemap* to compute the proportion of species belonging to each one of the 11 IUCN categories for each node of the taxonomy (see Supplementary Material for full details on the data and method used). These proportions were plotted with *pylifemap* using donut charts. The code leading to this visualization is given below; a snapshot is shown in Figure 1A.

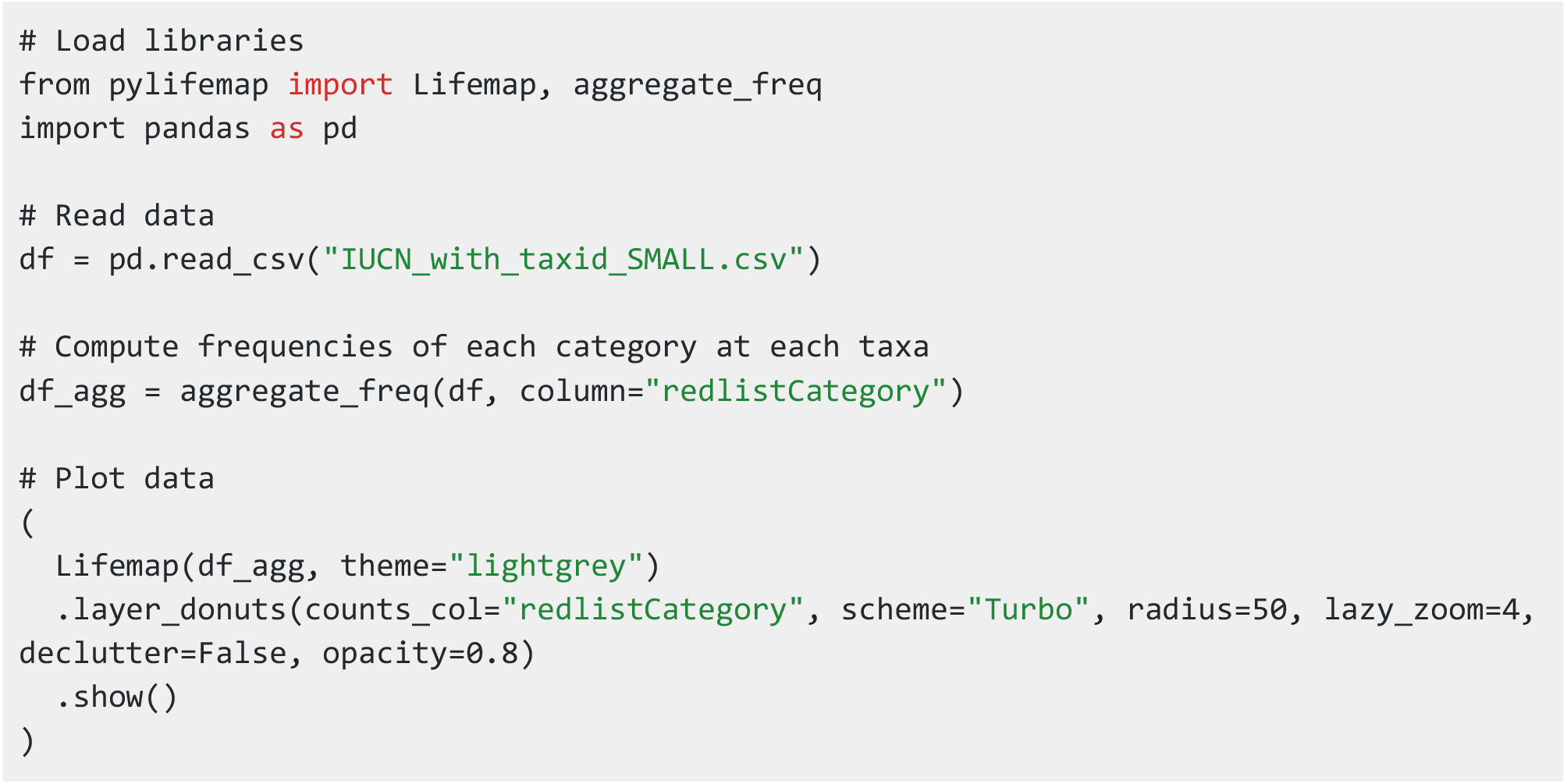

The example code illustrates some of the many visualization options proposed (listed in Table 1): the basemap theme can be chosen, the color palette (scheme) for categories can be changed, lazy loading of data is possible (i.e. only data associated with taxa visible at the current zoom level and around is displayed), decluttering (preventing elements overlap) can be activated or not, etc.

The obtained visualization (Figure 1A and online gallery) displays a clear view of the proportion of species with each IUCN category for each node of the taxonomy. Through an exploration of the map we can easily identify, for instance, that within Batrachia, the proportion of Caudata (comprising salamanders) being endangered or critically endangered (almost 50%) is much higher than within its sister group Anura (comprising frogs, around 15%, Figure 1A). This type of visualization also illustrates an important aspect of data visualization with *pylifemap*: even a large dataset of hundreds of thousands of entries can be smoothly explored, visualized, and ultimately interpreted because data are organized and viewed in a hierarchical manner, simply following the hierarchical nature of the entities (here taxa) to which they belong.

### Huanan seafood market metagenomics

Metagenomics studies involve the collection of diverse biological samples from a variety of environments, such as gut, soil, water, or air to gain insights into the microbial communities present in these ecosystems. Tools such as Kraken2 (Wood et al. 2019), STAT (Katz et al. 2021), or MetaPhlAn (Blanco-Míguez et al. 2023) are commonly utilized to identify and classify the taxonomic composition of the microbial communities present in these samples, using a reference database for the assignment. This taxonomic characterization is then used for downstream analyses, such as microbial diversity analyses, comparative analyses, and ecological modeling.

Visualization of taxonomic content data is critical in these analyses, and multiple tools have been developed for this purpose over the years (McMurdie and Holmes 2015; Wagner et al. 2018; Thrash et al. 2019; Breitwieser and Salzberg 2020). In all cases, the taxonomic classification viewers only display taxa on which reads have been mapped, and/or their ancestors, hindering a full picture of where reads did or did not map in the whole taxonomy. *pylifemap*, because it relies on the full NCBI explorer Lifemap, and also because multiple layers can be superimposed, overcomes this limitation and opens new perspectives.

The metagenomics data we used here are taxonomic assignations (performed with Kraken2) of environmental samples collected in the Huanan seafood market (Wuhan, China) in early 2020, a few days after the start of the SARS-CoV-2 pandemic (Liu et al. 2024). To each identified taxid is associated a number of reads uniquely pointing to it (referred to as “fragment_number”). Following (Crits-Christoph et al. 2024), and in order to best illustrate the capacities of *pylifemap* for complex data exploration, we decided to cross this metagenomics data with the list of 38 wild animal species that were found to be sold in the same market between May 2017 and November 2019 (Xiao et al. 2021), just before the beginning of the pandemic. Because these animals all belong to Amniota, we decided to subsample the metagenomics data to only the reads mapping to this clade.

Using the script below (see Supplementary Material for more explanation on file names), we used *pylifemap* to visualize these data using three layers (Figure 1B and online gallery): a point layer with size and color representing the “fragment number” from the Kraken2 output, a line layer to highlight the taxa present in the reference database used for the taxonomic assignation, and finally an icon layer to highlight – with a black cross – the wild species sold in the market (Xiao et al. 2021).

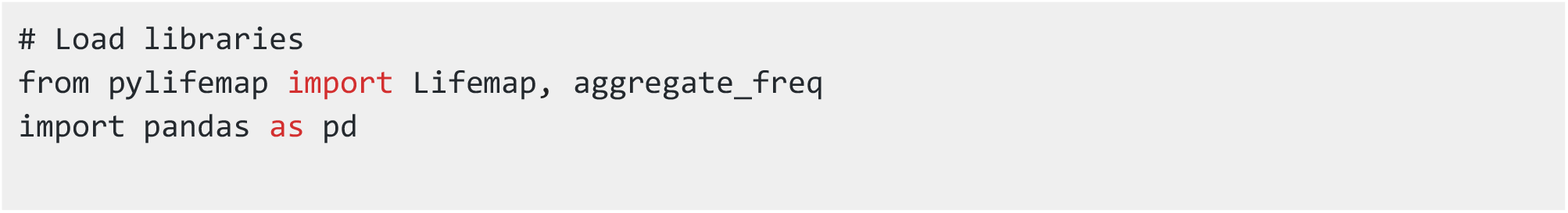
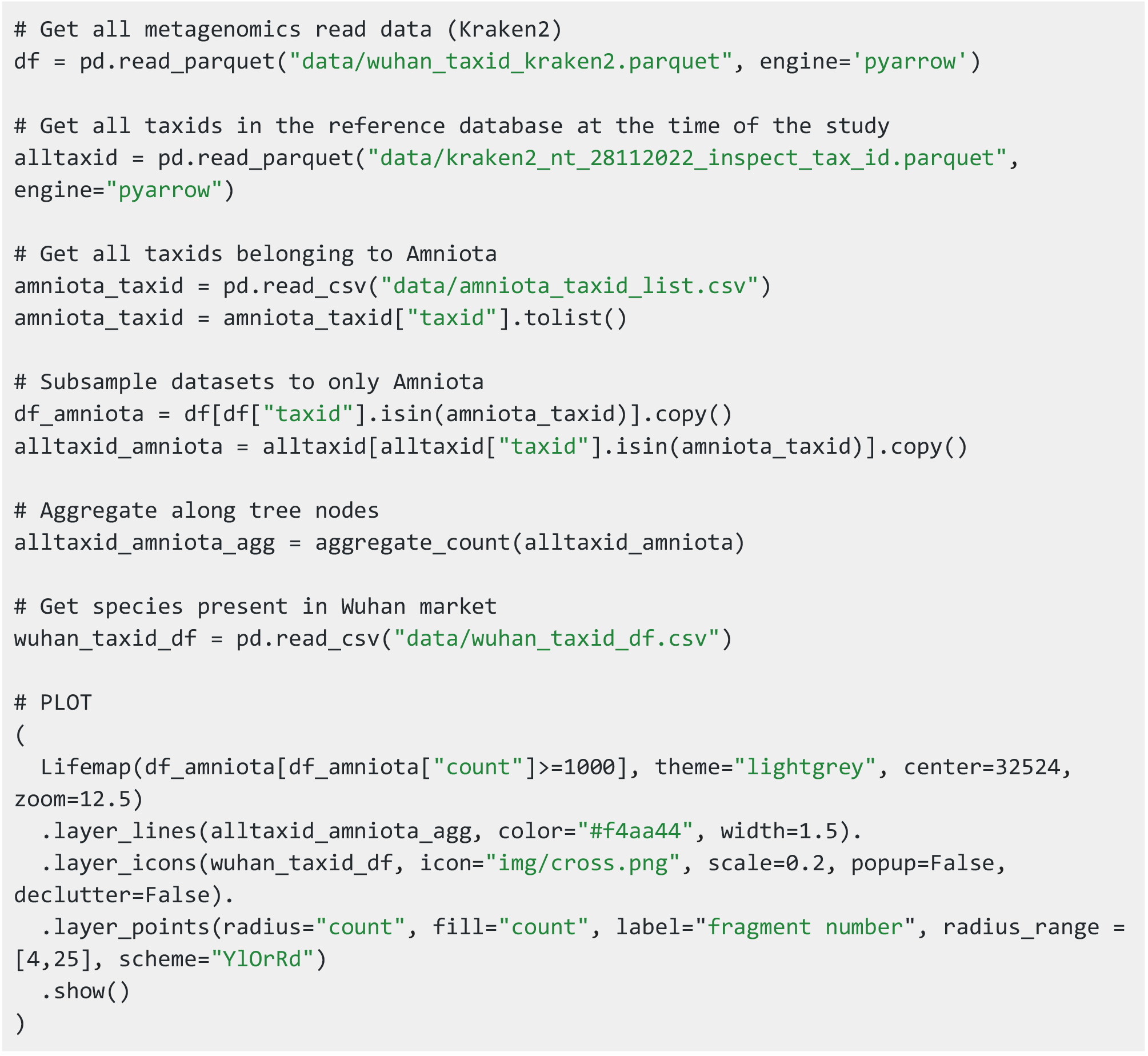

Interactive exploration of the obtained map allows an easy identification of taxa with large numbers of reads mapping to them. Looking at the black crosses, we also observe that some of these taxa correspond to wild species sold in the market prior to the pandemic. This is the case of the common pig (*Sus scrofa*), the Chinese bamboo (*Rhizomys sinensis*), Reeves’ muntjac (*Muntiacus reevesi*), Amur hedgehog (*Erinaceus amurensis*) and the raccoon dog (*Nyctereutes procyonoides*). Coloring the evolutionary path from the root to the list of species included in the reference database used by Kraken2 highlights interesting cases where the genomes of animals present in the market are not available in the reference database (like the masked palm civet *Paguma larvata*) or more complex cases where the taxonomic assignment occurs with known similar genomes (Siberian weasel, *Mustela sibirica*). The interactive visualization, taxonomy-oriented, also reveals thousands of reads assigned to species belonging to *Cetacea, Chiroptera, Gekkota*, which either suggests Kraken2 taxonomic assignment errors, or indicates the presence of living organisms found in the market (e.g. *Pipistrellus pipistrellus*) or species not reported as sold by the vendors (e.g. a turtle of the sub-order *Cryptodira*).

## Discussion and conclusion

The visualization tool we propose here with *pylifemap* consists in mapping data on a basemap representing the taxonomic relationships between all taxa present in the NCBI taxonomy.

This new visualization method for taxon-level biological data offers three main benefits. First, scalability: it allows users to explore very large datasets – up to millions of taxa and associated data – through an interactive map, making complex data easily navigable. Second comprehensiveness: it provides a complete taxonomic context by visualizing the entire taxonomy, rather than limiting the view to taxa with available data, as is common in other tools (e.g. Ondov et al. 2011; Breitwieser and Salzberg 2020). Finally, it offers an evolutionary perspective of the data at hand, essential for understanding biological patterns (“Nothing in biology makes sense except in the light of evolution”, Dobzhansky 1973).

This way of displaying and analyzing information directly on a map recalls Geographic Information Systems (GIS), i.e. computer systems where geographic data (location information) are linked with various descriptive information, enabling the visualization but also the analysis of data related to spatial relationships and patterns (for a review on GIS, see Wieczorek and Delmerico 2009). What we propose here could be referred to as a Taxonomic Information System (TIS), although very simple, where taxonomic data (the identity and localization of taxa on Lifemap) – in place of geographic data – are connected to various descriptive information and can be visualized and analyzed. This new way of treating biological data in a taxonomic context (i.e. as a TIS) may open new perspectives for the analysis and visualization of large taxonomy-centered datasets. Still, clear differences remain. For instance, GIS manipulates a basemap that is continuous and complete, when taxonomy might be incomplete or difficult to map to datasets that do not rely on taxonomy identifiers from the start, as seen with the two examples presented here. Similarly, spatial distances have a clear and constant meaning in GIS, when taxonomic distance is discrete, changes through time, and does not correlate with the distance observed in the Lifemap basemap. These differences show the limits of treating TIS as GIS, but drawing the parallel could undoubtedly inspire analytical principles to enrich the exploration of large taxonomy-structured datasets.

In conclusion, the *pylifemap* package provides an effective and user-friendly solution for visualizing and sharing taxon-level biological data on the NCBI taxonomic tree viewer, Lifemap. The package is highly customizable, allowing for easy computation and visualization of summary statistics at parent nodes. The demonstration of the utility of *pylifemap* on two contrasting biological datasets emphasizes its potential to explore and analyze large, complex and diverse datasets, making it a valuable resource for researchers in (meta)genomics, microbiology, ecology, and related fields, but also for outreach activities around taxonomy, systematics, and the Tree of Life.

## Software availability

The *pylifemap* Python package is distributed under the MIT license and available in the Python package index PyPI: https://pypi.org/project/pylifemap/. The source code is maintained on GitHub: https://github.com/Lifemap-ToL/pylifemap/. Extensive documentation, usage examples, and a gallery are available on the project website: https://lifemap-tol.github.io/pylifemap/. The current manuscript describes version 0.2.0 of *pylifemap*.

## Acknowledgment

This work was performed using the computing facilities of the CC LBBE/PRABI.

