## Supplementary Material and Methods for "pylifemap: Mapping Data onto the Tree of Life" (from another version of this preprint: version 1, 2025-12-16)

### 1. IUCN dataset preparation

Data download: the IUCN lists of species for each IUCN category were downloaded from the IUCN website [https://www.iucnredlist.org/](https://www.iucnredlist.org/) the 4th of December 2025 and stored in a csv file named `IUCN_all_assessments.csv`. This dataset cannot be redistributed here due to licensing restrictions but can be downloaded and built again from the official website.

The following python code was then used to get the NCBI taxonomy id (taxid) for each species in the IUCN list and save it to file `IUCN_with_taxid_SMALL.csv`. Taxonomy manipulations were performed with the *ete-ncbiquery* utility in the ete3 python package (Huerta-Cepas et al. 2016).

```
from ete3 import NCBITaxa
import pandas as pd

# Read input file
input_file = "IUCN_all_assessments.csv"
df = pd.read_csv(input_file, low_memory=False)

ncbi = NCBITaxa()
# OPTIONAL: ncbi.update_taxonomy_database()

# Function to get taxid
def get_taxid(name):
    try:
        taxid_list = ncbi.get_name_translator([name])
        return taxid_list.get(name, [None])[0]
    except Exception:
        return None

# Apply function to the column
df['taxid'] = df['scientificName'].apply(get_taxid)
df['taxid'] = df['taxid'].astype('string')

# Subsample
dfsmall = df[df['taxid'].notna()][['taxid', 'scientificName',
'redlistCategory','redlistCategoryCode']]

# Optional: save results
dfsmall.to_csv("IUCN_with_taxid_SMALL.csv", index=False)
```

### 2. Metagenomics dataset preparation

Metagenomics data from Liu et al. (2024) were retrieved from the Genome Sequence Archive (GSA) online (accession CRA010170). Kraken2 (Wood et al. 2019) was used to classify all reads, building the reference database from the Genebank nt version 28112022. Kraken2 outputs were stored in the file `wuhan_taxid_kraken2.parquet`. The list of taxids present in the *effective* Kraken2 database, i.e. the database effectively used by Kraken2 after the build process, was obtained with kraken-inspect and exported to `kraken2_nt_28112022_inspect_tax_id.parquet`.

Finally, the script below was used (i) to recover all taxids corresponding to taxa in the Amniota clade, and store them in the file `amniota_taxid_list.csv`, and (ii) to retrieve the taxids of the wild species sold in the market by (Xiao et al. 2021). Of note, *Corvus pectoralis* was changed to its synonymic name *Corvus pectoralis* before fetching the taxid numbers, and one species, *Arctonyx albogularis*, not found in the NCBI taxonomy database, was discarded for subsequent analyses. As before, all taxonomic operations were done with the *ete-ncbiquery* utility in the ete3 python package (Huerta-Cepas et al. 2016). All datasets used for this experiment are available from a dedicated data folder in the github repository of *pylifemap* at [https://github.com/Lifemap-ToL/pylifemap/tree/main/ doc/gallery/sources/data](https://github.com/Lifemap-ToL/pylifemap/tree/main/doc/gallery/sources/data).

```
from ete3 import NCBITaxa
import pandas as pd

# Get all taxid for all Amniota in NCBI taxononmy, and save to file
ncbi = NCBITaxa()
amniota_taxid = ncbi.get_descendant_taxa('Amniota')
amniota_taxid = pd.DataFrame(amniota_taxid, columns=["taxid"])
amniota_taxid.to_csv("amniota_taxid_list.csv", index=False)

# Get all taxid for wild species in Wuhan market
wuhan_latin_names = [
    "Nyctereutes procyonoides",
    "Erinaceus amurensis",
    "Mustela sibirica",
    "Arctonyx albogularis",
    "Meles leucurus",
    "Lepus sinensis",
    "Callosciurus erythraeus",
    "Paguma larvata",
    "Rhizomys sinensis",
    "Hystrix brachyura",
    "Muntiacus reevesi",
    "Myocastor coypus",
    "Marmota himalayana",
    "Vulpes vulpes",
    "Neovison vison",
```

```
    "Sciurus vulgaris",
    "Sus scrofa",
    "Trogopterus xanthipes",
    "Corvus pectoralis",
    "Spilopelia chinensis",
    "Pica pica",
    "Acridotheres cristatellus",
    "Alectoris chukar",
    "Phasianus colchicus",
    "Pavo cristatus",
    "Numida meleagris",
    "Orthriophis taeniurus",
    "Dinodon rufozonatum",
    "Bungarus multicinctus",
    "Sinonatrix annularis",
    "Gloydius brevicaudus",
    "Naja atra",
    "Naja kaouthia",
    "Ptyas mucosa",
    "Deinagkistrodon acutus",
    "Crocodylus siamensis",
    "Ptyas dhumnades",
    "Elaphe carinata"
]

# Get taxids
taxid_wuhan_dict = ncbi.get_name_translator(wuhan_latin_names)

# Create dataframe: missing names get NaN
df_wuhan_taxids = pd.DataFrame({
    "latin_name": wuhan_latin_names,
    "taxid": [taxid_wuhan_dict.get(name, [None])[0] for name in wuhan_latin_names]
})

# Save dataframe to parquet file
df_wuhan_taxids.to_csv("wuhan_taxid_df.csv", index=False)
```
